# Direct Observation Of Vesicle Transport On The Synaptic Ribbon Provides Evidence That Vesicles Are Mobilized And Prepared Rapidly For Release

**DOI:** 10.1101/2020.02.05.936153

**Authors:** Christina Joselevitch, David Zenisek

**Affiliations:** Department of Cellular and Molecular Physiology, Yale University School of Medicine, 333 Cedar Street, New Haven, CT 06511; Department of Ophthalmology and Visual Sciences, Yale University School of Medicine, 333 Cedar Street, New Haven, CT 06511; Department of Neuroscience, Yale University School of Medicine, 333 Cedar Street, New Haven, CT 06511; Department of Experimental Psychology, Psychology Institute, University of São Paulo, Av. Prof. Mello Moraes 1721, 05508-030 São Paulo-SP, Brazil

**Keywords:** synaptic depression, vesicle release, vesicle replenishment, vesicle transport, ribbon synapses, bipolar cells, retina, live cell imaging, TIRF microscopy, vesicle priming

## Abstract

Synaptic ribbons are thought to provide vesicles for continuous synaptic transmission in some retinal non-spiking neurons, yet recent studies indicate that genetic removal of the ribbon has little effect on vesicle release kinetics. To investigate vesicle replenishment at synaptic ribbons, we imaged synaptic vesicles and ribbons in retinal bipolar cells with TIRF microscopy during stimulation with trains of 30-ms depolarizations. Analysis of vesicles released by the stimuli revealed that the vast majority of releasable vesicles reside within 300 nm of the ribbon center. A single 30-ms step to 0 mV was sufficient to deplete the most membrane-proximal vesicle pool, while triggering rapid stepwise movements of distal vesicles along the ribbon and toward the plasma membrane.

Replenishment only becomes rate-limiting for recovery from paired-pulse depression for interstimulus intervals shorter than 250 ms. For longer interstimulus intervals, vesicle movement down the ribbon is fast enough to replenish released vesicles, but newly arrived vesicles are not release-ready. Notably, vesicle re-supply is 40-to 50-fold faster than previously measured in non-ribbon conventional synapses, whereas vesicle maturation rate is comparable. Moreover, in contrast to conventional synapses, vesicles docked at the base of the ribbon release with high fidelity. Lastly, our data show that with multiple stimuli, the delay in vesicle departure increases. Our results support a role for ribbons in the rapid supply and efficient preparation of vesicles for release, provide direct measurements of vesicle movement down the synaptic ribbon and suggest that multiple factors contribute to paired-pulse depression.

## INTRODUCTION

In non-spiking cells of the retina, cochlea and lateral line, synaptic ribbons harbor a dense array of glutamatergic synaptic vesicles near the plasma membrane directly opposite post-synaptic receptors. Because synaptic ribbons reside in neurons that release for prolonged periods of time, it has long been suggested that ribbon transport provides vesicles for uninterrupted continuous release [for reviews, see (Matthews and Fuchs, 2010; Moser et al., 2019)]. However, more recently this idea has been challenged.

Ribeye is the most abundant protein in synaptic ribbons and frame-shifting mutations to both *ribeye* genes in zebrafish leads to morphological changes and mislocalization of ribbons in lateral line hair cells, but minimal effects on exocytosis, even in response to very long stimuli (Lv et al., 2016). Similarly, genetic disruption of Ribeye in mouse leads to loss of all synaptic ribbons without change in the kinetics of release in bipolar cells (Maxeiner et al., 2016), or in the kinetics or amount of release in inner hair cells (Becker et al., 2018; Jean et al., 2018).

Moreover, despite the abundance of vesicles near ribbon sites, retinal ribbon-type synapses (but not all hair cell synapses (Goutman and Glowatzki, 2011; Levic et al., 2011) exhibit profound depression in response to pairs of stimuli, suggesting that ribbons may not be especially adept at resupplying vesicles (Burrone and Lagnado, 2000; Choi et al., 2008; Gomis et al., 1999; Mennerick and Matthews, 1996; Rabl et al., 2006; Sakaba et al., 1997; Singer and Diamond, 2003; von Gersdorff and Matthews, 1997).

If indeed synaptic ribbons mark the location of exocytosis (Midorikawa et al., 2007; Zenisek, 2008) and are responsible for the turnover of synaptic vesicles (LoGiudice et al., 2008), the time course of recovery from paired-pulse depression could reflect slow replenishment of release sites, slow maturation of newly arrived ribbon-attached vesicles, or both.

We set out to investigate whether vesicles are indeed moved down ribbons in response to stimuli and if so, the properties of this transport process. In addition, we investigated which mechanism, depletion or maturation, underlies the long recovery from synaptic depression in goldfish retinal bipolar cells. By directly imaging released and docked vesicles at synaptic ribbon sites with total internal reflection fluorescence (TIRF) microscopy, we found that vesicles on ribbons move toward the membrane in a rapid step-like manner and that vesicle replenishment only becomes rate-limiting for recovery from paired-pulse depression if interstimulus intervals are shorter than 250 ms. For longer interstimulus intervals, vesicle movement down the ribbon is fast enough to replenish released vesicles, but vesicles that newly arrive are not competent for release, indicating that both vesicle absence and biochemical steps downstream of vesicle arrival contribute to depression.

These observations are in stark contrast with those on conventional synapses, which report much longer resupply times (Midorikawa and Sakaba, 2015, 2017). Together, our results point to ribbons as both conveyor belts and priming devices, rendering ribbon-type synapses more efficient for continuous release than conventional ones.

## RESULTS

To investigate the properties of vesicle transport in bipolar cell synaptic terminals, synaptic vesicles in freshly dissociated bipolar cells from the goldfish retina were labeled with FM1-43, which loads into vesicles by endocytosis (Betz et al., 1992), and imaged using TIRF microscopy. Since vesicles are packed into the synaptic terminal at a higher density than can be resolved using light microscopy, labeling was restricted to a small fraction of the total vesicle pool in order to visualize individual synaptic vesicles (Zenisek et al., 2000). This was achieved by exposing cells briefly to FM1-43 in the presence of 25 mM K^+^ (see *Methods*). After loading with the dye, bipolar cells were voltage-clamped at the cell soma. In many experiments, a Ribeye-binding peptide conjugated with rhodamine was added to the patch pipette (Joselevitch and Zenisek, 2009; Zenisek et al., 2004), allowing us to visualize ribbon sites in the same cells.

### Immobilized Synaptic Vesicles Gather at Ribbon Sites

Synaptic vesicles were visible as diffraction-limited spots after labeling. As described previously (Holt et al., 2004; Zenisek, 2008; Zenisek et al., 2000), most vesicles fluctuate in intensity with time and were visible for less than 200 ms, whereas a subset of vesicles retained a steady fluorescence intensity, indicating that they were immobile. When bipolar cells were periodically subjected to 30-ms step depolarizations from −60 to 0 mV, fusion of immobilized vesicles could be seen as an increase in spot fluorescence followed by the formation of a cloud of dye, as reported elsewhere (Zenisek et al., 2000). An example of such an event is illustrated in **Figure 1A-B** and in **supplemental movies 1 and 2.**

**Figure 1.**
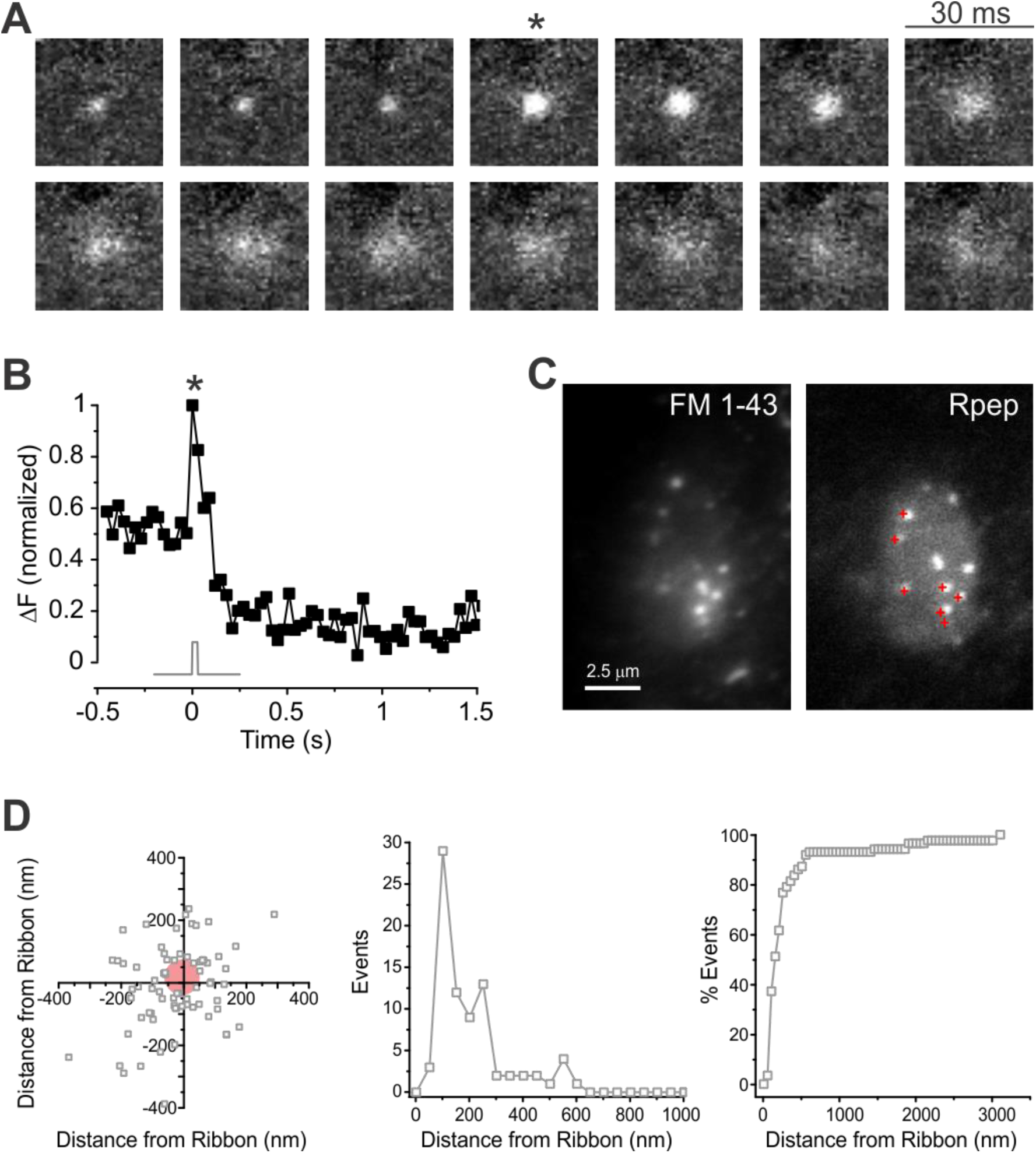
Released vesicles concentrate at ribbon sites. (A) Consecutive video images from a docked vesicle that underwent fusion. Released vesicles appear as a cloud of dye and may be visible at the cell membrane for a long period of time. The asterisk indicates the frame of a single 30-ms depolarization from −60 to 0 mV. (B) The normalized fluorescence of the vesicle in (A) is plotted in relation to the timing of the voltage stimulus (grey trace). The release of vesicular contents is seen as an increase in fluorescence and correlates well to the change in membrane potential. The asterisk corresponds to the timing of depolarization in (A). (C) Release events in a bipolar cell terminal seen with FM1-43 (left) correlate well with ribbons labeled in the same terminal by the Ribeye-binding peptide (*Rpep*, right). Released vesicles are marked by red crosses in the Rpep image. The FM image is an average of 85 frames and the Rpep image is an average of 10 frames. (D) High-resolution scatter plot (left), simple histogram (center) and cumulative histogram (right) of 87 events observed in 25 records from 9 bipolar cells, showing that most vesicles docked within 300 nm of labeled ribbons. These active release zones correspond roughly to the base of synaptic ribbons. Red circle indicates region devoid of vesicles near the center of the ribbon.

Previous reports indicated that docked vesicles preferentially localize near ribbons (Vaithianathan et al., 2016; Zenisek, 2008) and that brief stimuli give rise to fusion events mostly near active zones (Zenisek et al., 2000) or labeled ribbon sites (Midorikawa et al., 2007), whereas longer depolarizations elicit extrasynaptic and non-vesicular fusion events (Coggins et al., 2007; Midorikawa et al., 2007; Zenisek et al., 2000). Like these previous reports, we found that vesicle docking and exocytosis colocalized well with ribbon sites. **Figure 1C** shows a map of the location of fusing vesicles in one terminal (left panel and **supplemental movie 3**), marked as red crosses and superimposed on a TIRF image of the synaptic ribbon locations taken just after stimulating exocytosis (right panel). As can be observed in this example, fusion events were confined to the vicinity of ribbon locations.

To investigate the association between vesicles and ribbon in more detail, we mapped the distance for each vesicle relative to the nearest ribbon at high spatial resolution. For this, we localized the center of each fluorescent spot to sub-pixel accuracy, a strategy used to visualize small movements of molecular motors (Yildiz et al., 2003) and employed by super-resolution microscopy techniques (Toomre and Bewersdorf, 2010). Overall, 79% of vesicles that could be localized with an error equal to or less than 20 nm (see *Methods*) were found within 300 nm of the center of the nearest ribbon. The distribution of these vesicles relative to the ribbon center is shown as a scatter plot in the left panel of **Figure 1D** and replotted as a histogram of distances in the center panel and as a cumulative histogram in the right panel.

Since the base of a synaptic ribbon is around 400 nm long (von Gersdorff et al., 1996), we propose that each of these active release zones corresponds to the base of a synaptic ribbon. It is interesting to note that we found a void of approximately 50 nm radius in the center of the scatter plot (**Figure 1D**, left panel). This roughly corresponds to the width of the ribbon along its minor axis (von Gersdorff et al., 1996) and suggests that vesicles do not dock directly underneath synaptic ribbons, probably because this is prevented by the protein complexes involved in anchoring ribbons at the membrane (Dick et al., 2003; tom Dieck et al., 2005). Indeed, the region directly beneath the ribbon appears devoid of vesicles in electron micrographs of bipolar cells (von Gersdorff et al., 1996). Evidently, bipolar cell synaptic release is mostly confined to ribbon sites when triggered by brief stimuli, as previously proposed (Coggins and Zenisek, 2009; Midorikawa et al., 2007; von Gersdorff et al., 1996; Zenisek et al., 2000). In the following sections, we refer to active zones as these sites of concentrated synaptic release and pool data from both Ribeye-labeled cells, where direct correlation to ribbon sites is possible, and unlabeled cells, where the signal-to-noise ratio allows for better imaging of vesicles. The remaining analysis in this paper was restricted to this ribbon-associated pool of vesicles.

### Vesicles Move Down the Ribbon Upon Stimulation

Bipolar cells exhibit at least two kinetic components of exocytosis in response to step depolarizations to 0 mV (Mennerick and Matthews, 1996; Neves and Lagnado, 1999; Sakaba et al., 1997; Singer and Diamond, 2003; Zenisek et al., 2000). The fastest component of exocytosis is rate-limited by the activation of the Ca^2+^ current and depleted with a time constant of few milliseconds (Mennerick and Matthews, 1996; Sakaba et al., 1997; Singer and Diamond, 2003). The rapidity of exocytosis coupled with its insensitivity to Ca^2+^ (Heidelberger et al., 1994) or Ca^2+^ buffers (Mennerick and Matthews, 1996; Sakaba et al., 1997), and the tight colocalization of ribbon sites to Ca^2+^ channels (Zenisek et al., 2003) strongly suggest that this rapid component must arise from vesicles at the base of the ribbon. A second, slower component is depleted with a time constant of over 250 ms (Burrone and Lagnado, 2000; Mennerick and Matthews, 1996; Sakaba et al., 1997) and was suggested to arise from vesicles both at active zones and at outlier locations (Mehta et al., 2014; Midorikawa et al., 2007; Zenisek, 2008; Zenisek et al., 2000).

We therefore looked more directly at the possibility that vesicles move down the ribbon in response to short stimuli. Bipolar cells were depolarized from −60 to 0 mV for 30 ms to elicit exocytosis of the vesicles docked at the base of the ribbons and subsequent replenishment. Vesicle movement toward the coverslip, as indicated by an increase in vesicle fluorescence without the lateral spread of dye fluorescence, often followed these brief depolarizations (**Figure 2A, supplemental movie 4**).

**Figure 2.**
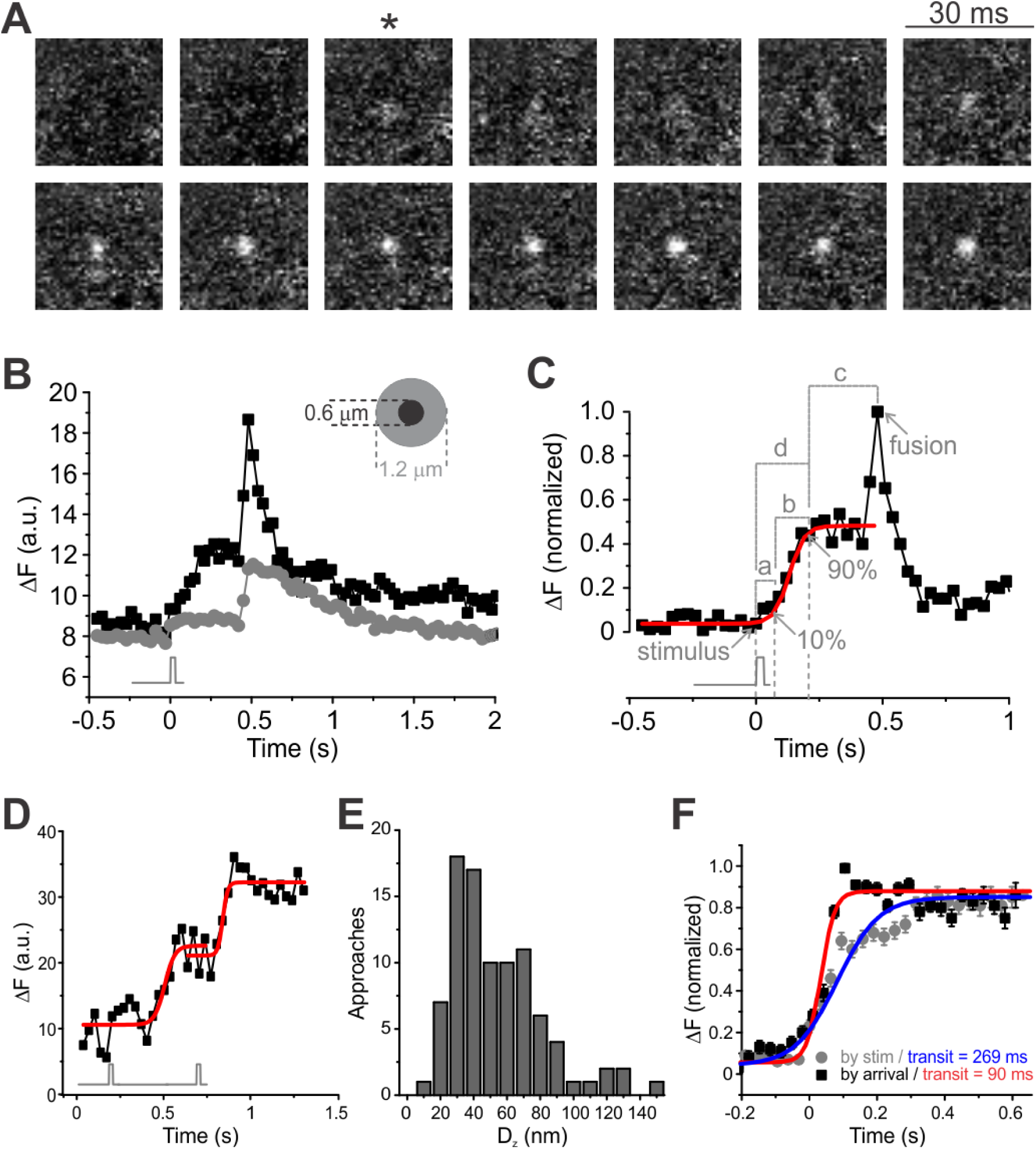
Vesicles move down the ribbon upon stimulation. (A) Voltage stimuli trigger capture and movement of vesicles move down the ribbon. Consecutive video images from a newly immobilized vesicle; the asterisk indicates the frame of a single 30-ms depolarization from −60 to 0 mV. (B) Quantifying triggered movement along the synaptic ribbon. The grey trace indicates the timing of a single 30-ms depolarization from −60 to 0 mV. The fluorescence of a newly immobilized vesicle (black symbols) increases as the vesicle approaches the membrane, and this increase is strictly correlated with stimulus timing. The fluorescence is confined to a 0.6-μm diameter circle (indicated in the inset); the fluorescence of the surrounding area (grey symbols) does not increase in a similar fashion. (C) The fluorescence profile of individual approaches (black squares) was fitted with a Boltzmann function (red trace) and the following parameters were measured for each vesicle: (*a*) *delay for departure*, defined as the difference between the *departure time* (10% of the maximal amplitude of the Boltzmann function) and the *stimulus time* (grey trace under graph); (*b*) *transit time*, defined as the difference between the *arrival time* (10% of the maximal amplitude of the Boltzmann function) and the *departure time*; (*c*) *sitting time*, defined as difference between the *fusion time* and the *arrival time;* and (*d*) *delay for arrival*, defined as the difference between the *arrival time* and the *stimulus time*. (D) Example of a vesicle that makes two consecutive steps towards the membrane. Each step could be well fit by a Boltzmann function (red traces) with 126.9 ms and 60.5 ms transit times, respectively. The black trace indicates stimulus timing. (E) Triggered vertical displacement along the ribbon. Distance traveled by each of 94 approaches within a single step towards the membrane, calculated from the ratio of fluorescence between the start and end of the movement (see *Methods*). (E) Mean fluorescence profile of 92 approaches aligned to the *arrival time* (black squares) or to the *stimulus time* (grey circles). Boltzmann fits to these functions (red and blue traces) yielded distinct *transit times*: 90 ms when vesicles were aligned to the *arrival time* and 269 ms when vesicles were aligned to the *stimulus time*. This means that vesicles are able to reach the membrane within 90 ms but start moving with a variable delay after the cell is depolarized. Error bars are S.E.M.

**Figure 2B** plots the fluorescence profile of the vesicle in **Figure 2A** (black symbols) and of a contiguous annulus (grey symbols), used to estimate background fluorescence (details of this analysis are in *Methods*). The timing of vesicle arrivals at active zones was determined as previously described (Zenisek et al., 2000) by fitting the fluorescence of approaching vesicles to a sigmoidal function (**Figure 2C, red trace**). The *departure time* was defined as when that fit reached 10% of its maximum and the *arrival time* as when it reached 90% of the asymptote. The time vesicles spent moving toward the membrane was calculated as the difference between the *departure time* and *arrival time* and was called *transit time* (**Figure 2C**, *b*). Other measured parameters were: *delay for departure* (**Figure 2C**, *a*), defined as the difference between the *departure time* and the *stimulus time*; *sitting time* (**Figure 2C**, *c*), or the difference between the *fusion time* and the *arrival time;* and *delay for arrival* (**Figure 2C**, *d*), defined as the difference between the *arrival time* and the *stimulus time*.

The movement of approaching vesicles at active zones was directionally biased toward the membrane, since 92 increases in vesicle fluorescence were observed in the frames following a depolarization and no stepwise decreases in fluorescence were observed (39 movies, 12 cells). Occasionally, vesicle brightening events were observed twice in the same location, suggesting a vesicle making two steps towards the membrane (*n* = 17 vesicles, 11 movies, 7 cells). An example is shown in **Figure 2D**. Each approach could be fit by the same sigmoidal function, yielding similar individual *transit times* (115.5 ± 46.2 ms after the stimulus for the first approach and 151.2 ± 46.3 ms for the second, mean ± S.E.M.), consistent with stepwise movements towards the cell membrane.

**Figure 2E** shows a histogram of the measured distances traveled by each of 94 stepwise approaches analyzed with the method in **Figure 2C**. It is noteworthy that the histogram peaks at 30-40 nm (median = 44.5 nm, mean ± SD = 54.4 ± 39.4), a vertical displacement similar to the distance measured between vesicles on the ribbon (von Gersdorff et al., 1996). **Figure 2F** shows the average fluorescence of 92 such approaches aligned to the voltage stimulus (grey circles) or to the *arrival time* (black squares). When aligned to the *arrival time,* the resulting mean intensity profile could be fit by a sigmoidal function with an apparent *transit time* of 90 ms. When the same data was aligned to the triggering stimulus, however, the resulting curve is broader, with an apparently longer *transit time* (269 ms), indicating that even though approaches tend to be fast, not all vesicles start moving at the same time after a stimulus.

### Some Vesicles Start Moving Later

We next analyzed the distribution of *transit times* (**Figure 3A**), *delays for departure* (**Figure 3B**) and *delays for arrival* (**Figure 3C**) in the first 240 ms following a depolarization. **Figure 3A** shows that although 70% of observed vesicles made their way to the membrane within 150 ms, the distribution of *transit times* exhibited variability, which can also be seen in the distribution of *delays for departure* (**Figure 3B)**. The coefficient of variation of the latter distribution (defined as the mean divided by the SD) was larger: *cv* = 1.1 for *transit times* and 1.54 for *delays for departure.* As shown in **Figure 3D**, both *transit times* and *delays for departure* contribute to *delays for arrival*, because the latter correlate with both variables, the correlation with *transit times* being higher (adjusted *R*^2^ = 0.66 for *transit times* and 0.43 for *delays for departure,* respectively).

**Figure 3.**
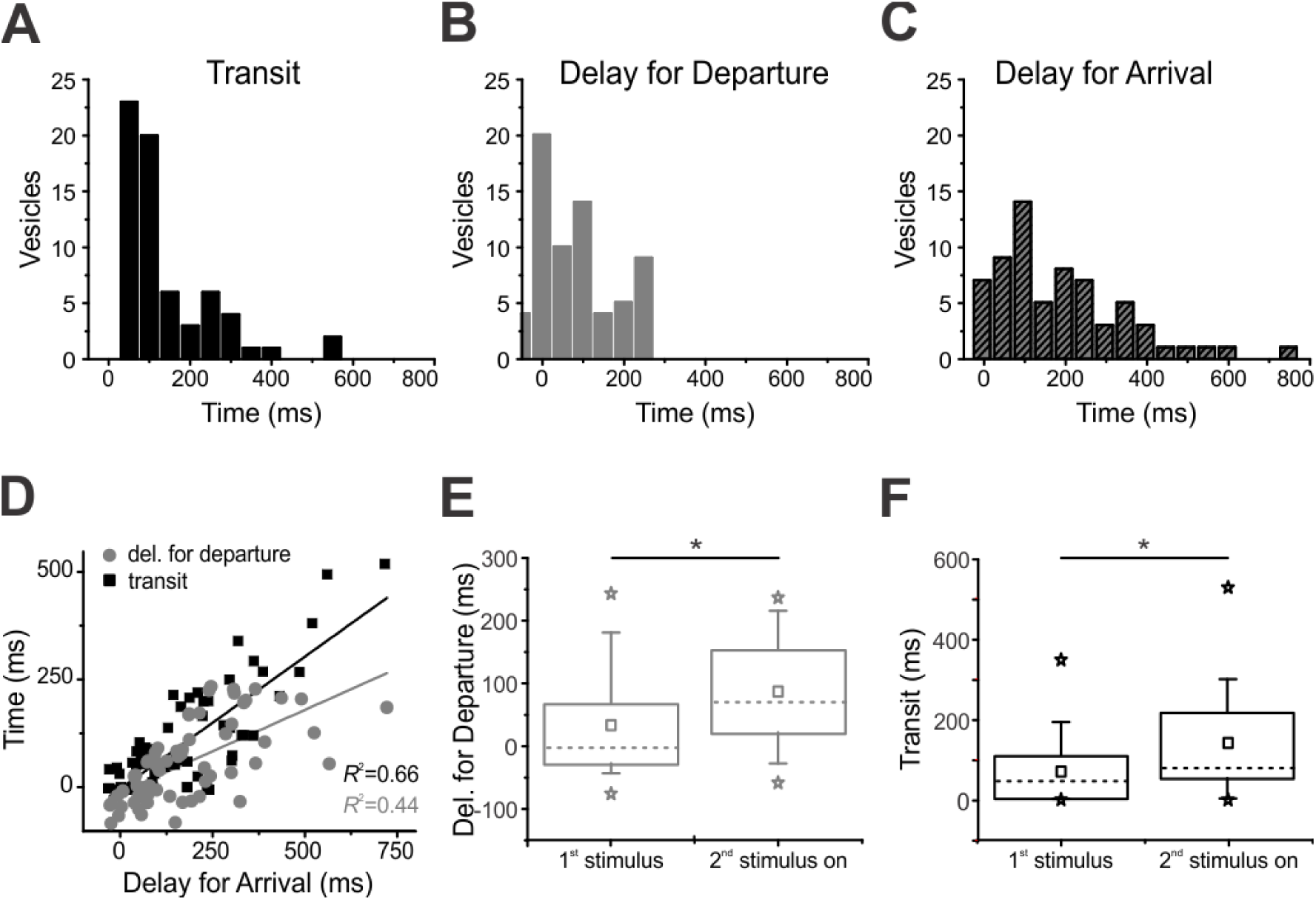
Arrival depends on how long vesicles travel and when vesicles depart. (A) Distribution of *transit times* in 66 vesicles (12 cells, 12 movies). 74.2% of observed vesicles arrived at the membrane within 150 ms. (B) Distribution of *delays for departure* for the same vesicles in (A). The variability in *delays for departure* is larger than that of *transit times (coefficient of variation* = 1.1 for *transit times* and 1.54 for *delays for departure*). (C) Distribution of *delays for arrival* in the same vesicles of (A) and (B). (D) Both *transit times* (black squares) and *delays for departure* (grey circles) contribute to *delays for arrival*. Straight lines are linear regressions to the data (*R*^2^ = 0.66 and 0.44 respectively). (E) and (F) *Delays for departure* and *transit times* are influenced by stimulus history. Both median values (indicated in the figures by the dashed lines) are smaller for the first stimulus in comparison to the remainder applied in all trains. Means are shown as squares, the boxes enclose the interquartile range (between the 25^th^ and 75^th^ percentiles), the bars indicate the 10^th^ and 90^th^ percentiles, the stars are the minimum and maximum, and the asterisks denote level of significance (*P* = 0.011 for *delays for departure* and 0.010 for *transit times*, two-tailed Mann-Whitney test).

We next investigated whether the stimulus history influences *delays for departure* and *transit times*. We applied trains of 30 ms depolarizations from −60 mV to 0 mV with different interstimulus intervals (240, 250 and 480 ms) to deplete the bottom row of vesicles and elicit distinct degrees of replenishment, as reported elsewhere (Gomis et al., 1999; Mennerick and Matthews, 1996; Sakaba et al., 1997; von Gersdorff and Matthews, 1997). We then analyzed *delays for departure* (**Figure 3E**) and *transit times* (**Figure 3F**) for the first stimulus separately in relation to the remainder stimuli of each train.

As shown in **Figure 3E**, *delays for departure* were significantly shorter for the first stimulus (median = −2 ms for the first stimulus and 70 ms for the second stimulus, two-tailed Mann-Whitney *U* = 343, *n*_1_<>*n*_2,_ *P* = 0.011. The negative median reflects noise in the fluorescence data; some vesicles appear to start moving prior to the triggering stimulus. Similarly, **Figure 3F** shows that *transit times* were also shorter for vesicles departing after the first stimulus (median = 48 ms for the first stimulus and 81 ms for the second stimulus, two-tailed Mann-Whitney *U* = 342, *n*_1_<>*n*_2_, *P* = 0.010). This result suggests that vesicles wait longer to move and need longer to reach the membrane after a second stimulus than following a first stimulus.

### Estimate of Vesicle Speed on the Ribbon

We next set out to estimate how fast synaptic vesicles move down the ribbon. To do so, we used two different methods, illustrated in **Figure 4**. The first method consisted in fitting the mean fluorescence aligned by *arrival time* of the 92 approaches in **Figure 2F** (**Figure 4A**, black squares) to an exponential function followed by an abrupt stop (**Figure 4A**, grey circles). Using the time constant of the fit and the measured length constant of our evanescent field, we were able to estimate a speed of each vesicle in transit. This method yielded an estimate of 870 nm/s. Alternatively, we used the same model in **Figure 4A** to individually estimate the speed of 82 approaches (ten of the 92 approaches could not be fit). With this approach, the calculated speed was 595 ± 42 nm/s (mean ± S.E.M.; median = 497 nm/s); the distribution is shown in **Figure 4B**. **Figure 4C** shows that vesicle speed is also influenced by stimulus history -speeds were significantly higher for the first stimulus (median = 600 nm/s for the first stimulus and 405 nm/s for all subsequent stimuli, two-tailed Mann-Whitney *U* = 1090.5, *n*_1_<>*n*_2,_ *P* = 0.014).

**Figure 4.**
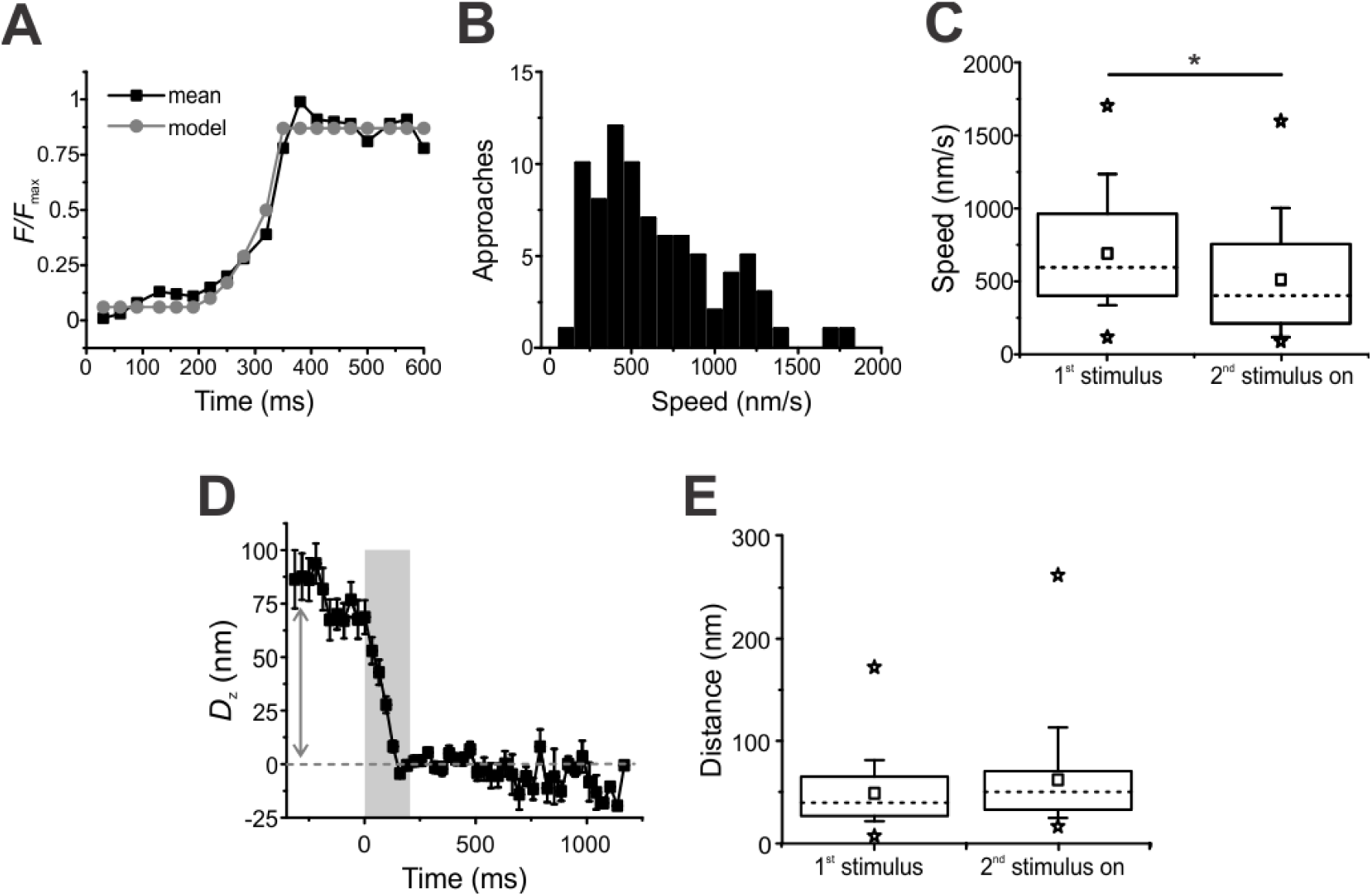
Methods for estimating vesicle speed. (A) The mean fluorescence profile of 92 approaches aligned to the *arrival time* (black squares, same as the black squares in Figure 2F) was modeled with an exponential rise followed by an abrupt stop (grey circles), yielding a speed estimate of 870 nm/s. (B) Distribution of the fluorescence of 82 individual approaches (12 cells, 36 movies) fit with the model in (A). Average speed was 595 ± 42 nm/s (mean ± S.E.M., median = 497 nm/s). (C) Stimulus history influences speed. Both medians (dashed lines) and means (squares) are smaller from the second stimulus on. Boxes enclose the interquartile range (between the 25^th^ and 75^th^ percentiles), the bars indicate the 10^th^ and 90^th^ percentiles, the stars are the minimum and maximum, and the asterisks denote level of significance (*P* = 0.014, two-tailed Mann-Whitney test). (D) Vertical displacement for the fluorescence of 92 approaches aligned to arrival time (black squares in Figure 2F). Error bars are S.E.M. The mean total displacement was 78.46 nm (arrow); dividing this number by the mean transit for this dataset (90 ms) yields a speed estimate of 871 nm/s. (E) There is no significant difference in distance traveled for first stimulus and subsequent stimuli. Although median (dashed lines) and mean (squares) speeds are larger from the second stimulus on, the difference is not statistically significant (*U =* 820.5, *P* = 0.13, two-tailed Mann-Whitney test). Boxes enclose the interquartile range (between the 25^th^ and 75^th^ percentiles), the bars indicate the 10^th^ and 90^th^ percentiles, the stars are the minimum and maximum.

Lastly, we also calculated speed from the vertical displacement (*D*_z_) of the same 92 approaches aligned to the *arrival time* (**Figure 2F**, black squares), based upon the relative fluorescence of the vesicle before and after its approach, and according to the formula described in *Methods*. **Figure 4D** depicts a plot of the mean vertical distance traveled by these vesicles towards the membrane, showing that they traveled the depth of our evanescent field within less than 250 ms (grey area). The mean vertical displacement in this graph is 78 nm (arrow); dividing this number by the mean transit of the vesicles aligned to arrival time (90 ms, red line in **Figure 2F**) yields a speed estimate of 871 nm/s. All three methods produced values close to the 800 nm/s estimate obtained elsewhere during a continuous stimulus (Zenisek et al., 2000), which let more Ca^2+^ into the cell than our brief depolarizations. This suggests that the extra Ca^2+^ during a prolonged stimulus does not make vesicles move faster.

Finally, we checked whether stimulus history influenced the distance each vesicle traveled following a stimulus. **Figure 4E** shows the results for the 90 of the 92 approaches in **Figure 2E**. Although the distances traveled from the second stimulus on were on average larger (mean ± S.E.M. = 48.5 ± 4.8 nm for first stimulus and 61.7 ± 6.6 nm for the second stimulus on), the difference did not reach statistical significance (medians = 39.3 nm and 49.9 nm for first and second stimulus on, respectively; two-tailed Mann-Whitney *U* = 820.2, *n*_1_<>*n*_2_, *P* = 0.013). To summarize, the results shown so far indicate that from the second stimulus on, vesicles depart later, take longer to reach the membrane, and travel slower.

### Triggered Vesicle Replenishment is Incomplete for Short Interstimulus Intervals

To investigate further how quickly vesicles are replenished, we compared the numbers of stimulus-elicited fusion events and newly arrived vesicles at active zones for trains of 30 ms steps from −60 mV to 0 mV at different interstimulus intervals (60, 120, 250 and 480 ms). The results obtained with the 250 ms interstimulus interval protocol are depicted in **Figure 5A-B**. **Figure 5A** shows a histogram illustrating when fusion events (black columns) and new vesicle arrivals (white columns) occurred in response to a series of four 30 ms depolarizations to 0 mV at 4 Hz. Fusion events were time-locked to the depolarizations, with no events occurring during the interval between stimuli. As previously described, the synapse exhibited depression (Gomis et al., 1999; Mennerick and Matthews, 1996; Sakaba et al., 1997; von Gersdorff and Matthews, 1997); the number of released vesicles decreased drastically from the first stimulus to the remainder in the train. **Figure 5B** plots the data in **Figure 5A** as a cumulative histogram. Of note, the number of recruited vesicles following a particular depolarization (white circles) mimicked the number of fusion events observed during the same stimulus (black squares), indicating little vesicle depletion during the stimulus protocol.

**Figure 5.**
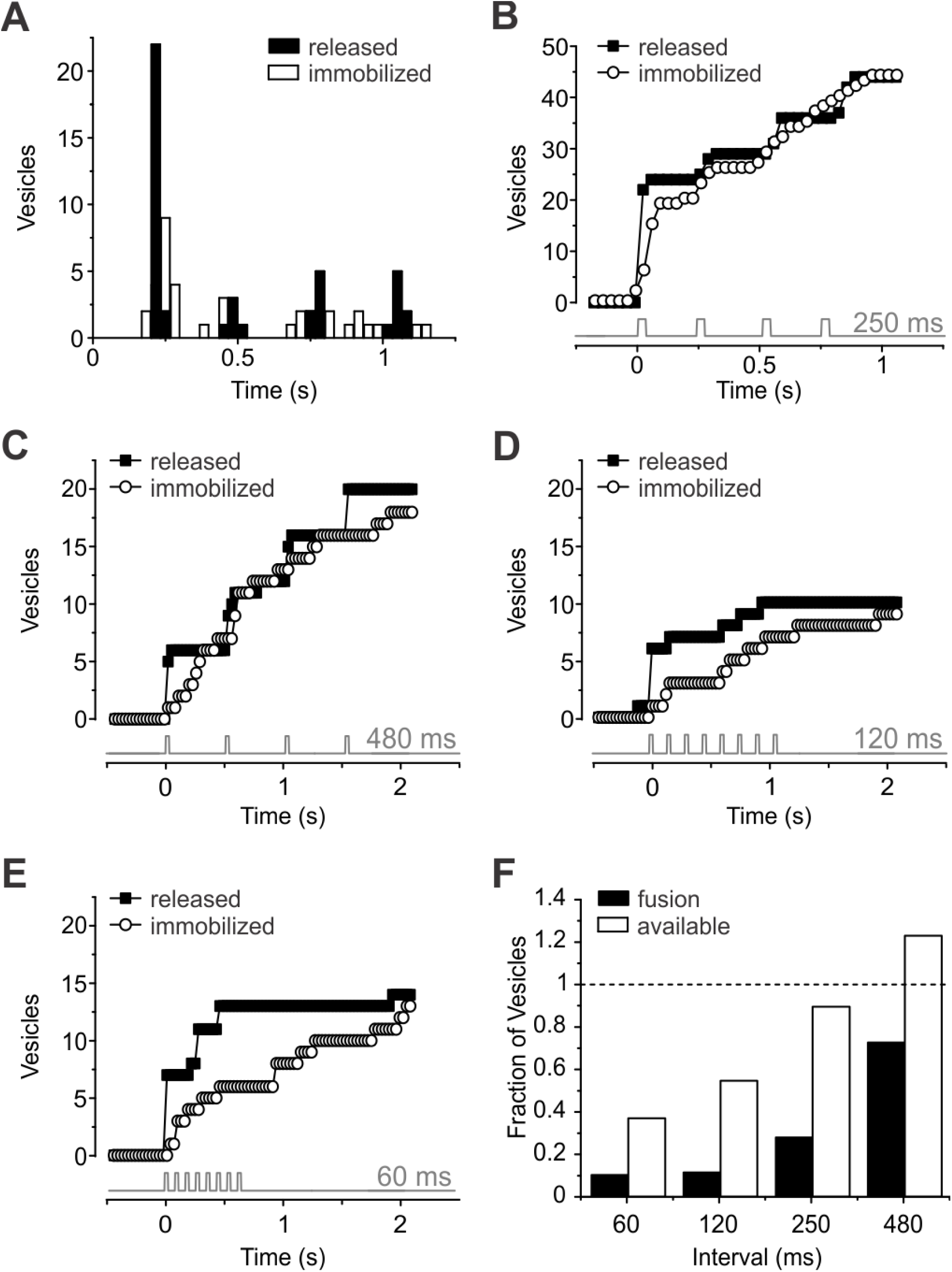
Timing of vesicle exocytosis and translocation in response to train of stimuli. (A) Histogram showing timing of fusion events (black columns) and capture events (white columns) in response to series of four 30-ms depolarizations from −60 to 0 mV at 4Hz. Timing of depolarizations is shown at the bottom of the graph in (B). Data from 4 cells, 19 movies. (B) Cumulative histogram of the data in (A). Fusion events are depicted as black squares, and captured vesicles as white circles; grey trace indicates the timing of each voltage step. During the trains, the total number of newly recruited vesicles follows closely the number of released vesicles. (C) Cumulative histogram of fusion events (black squares) and capture events (white circles) for the protocol with 480 ms interstimulus interval (*n* = 5 cells, 6 movies). Also here, the number of immobilized vesicles follows closely the number of released vesicles. (D) Cumulative histogram of fusion events (black squares) and capture events (white squares) for the protocol with 120 ms interstimulus interval (*n* = 5 cells, 5 movies). For this protocol, the number of immobilized vesicles falls short of the number of released vesicles at all times during the voltage train but catches up by the end of the movie. (E) Cumulative histogram of fusion events (black squares) and capture events (white squares) for the protocol with 60 ms interstimulus interval (*n* = 4 cells, 5 movies). Here, vesicle depletion is most pronounced. (F) Fractional vesicle occupancy and fusion events for all depolarizations after the first one in a train, normalized to the number of fusion events in the first depolarization. The occupancy at each depolarization (available) was estimated by taking the number of newcomer vesicles and subtracting the number of vesicles lost by exocytosis. Note that replenishment maintains an available pool of vesicles for longer intervals, but fails to fully replenish vesicles following shorter intervals (white bars).

**Figure 5C-E** shows the results obtained with three other interstimulus intervals: 480 ms (**Figure 5C**), 120 ms (**Figure 5D**), and 60 ms (**Figure 5E**). While there are clear signs of vesicle depletion at interstimulus intervals shorter than 250 ms (**Figure 5D-E**), for trains of lower frequency the number of newly immobilized vesicles followed closely the number of released vesicles (**Figure 5A-C**). **Figure 5F** shows the number of available vesicles and the number of released vesicles relative to step 1 for all subsequent steps at different train frequencies. To determine the number of available vesicles, the number of newly added newcomer vesicles were counted and the number lost via exocytosis were subtracted from that number for each step in a train after the initial step. The results show that ribbons facilitate nearly complete vesicle re-supply for intervals >250 ms between pulses, but fail to keep up when intervals are shorter. Therefore, a dearth of vesicles contributes to depression for these short interstimulus intervals. At longer intervals, however, vesicles are present, but still unable to undergo exocytosis. The time needed for fusion after recruitment, or *sitting time*, could play a role in this phenomenon.

### Newly Arrived Vesicles Are Not Fusion Competent

We next investigated how long it takes for a vesicle to fuse once it has arrived. To do so, we measured all vesicle arrival times using the sigmoidal fit approach highlighted above and measured the delay between each arrival time and subsequent stimuli. This delay was defined as the *sitting time*. For each sitting time, we determined which fraction of vesicles fused in response to a depolarization. **Figure 6A** shows the results for the 47 vesicles with highest signal-to-noise ratio in our dataset, binned for different delays (*n* = 119 events, 47vesicles, 36 movies, 13 cells).

**Figure 6.**
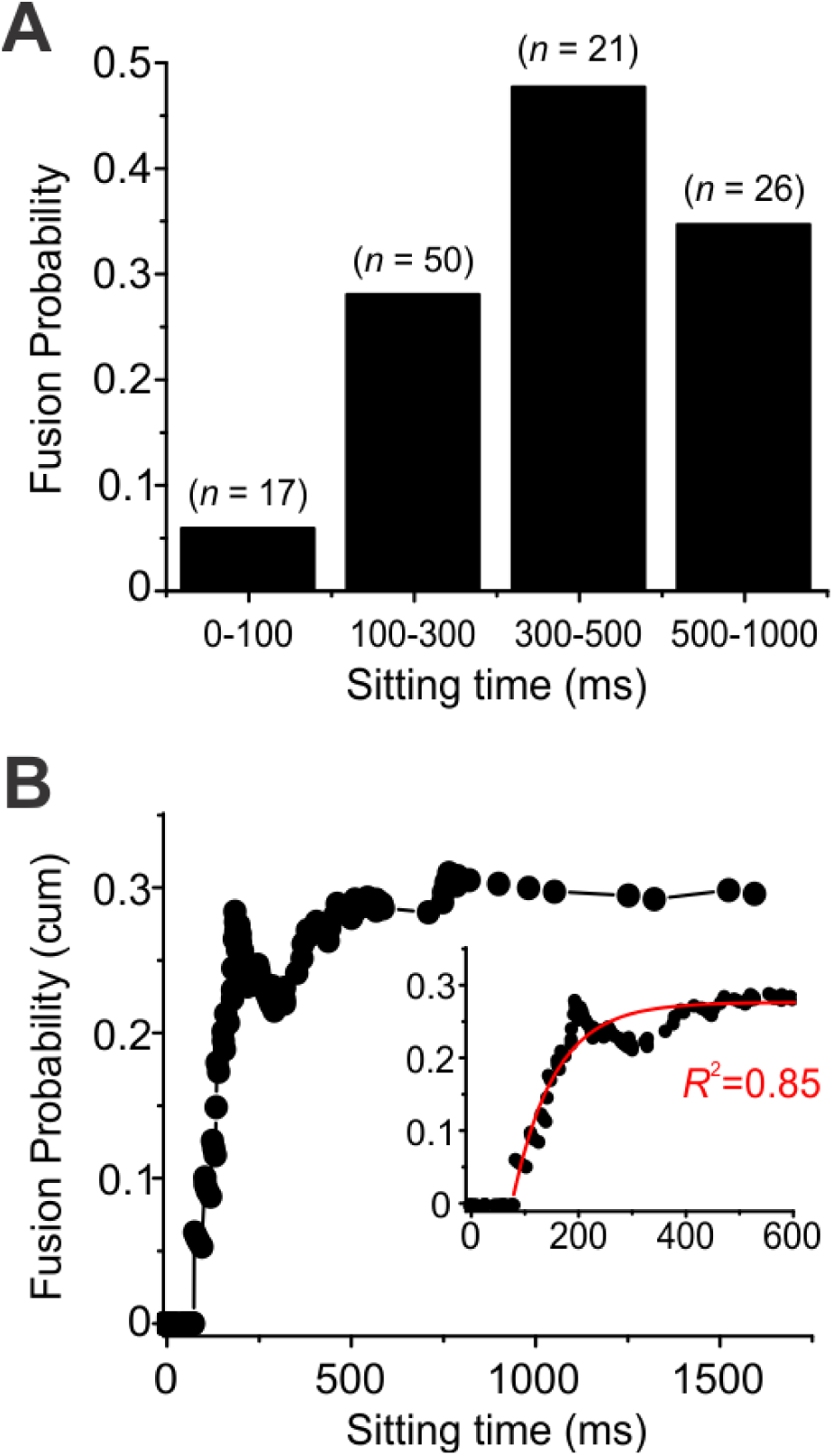
Newly arrived vesicles are not fusion competent. (A) Probability of vesicle fusion for different intervals between the time of vesicle arrival and fusion (*sitting time*). Note that vesicle fusion rate increased with sitting time (*n* = 119 events, 47 vesicles; 5 vesicles had sitting time > 1000 ms and are not included in the graph). (B) Same data set as in (A), but plotted as a cumulative probability of fusion for all vesicles with each sitting time or briefer. Within the first 90 ms after arrival at the membrane, the probability of fusion is zero. The graph could be fitted with a single exponential function with time constant of 76.87 ms (insert, *y*_0_ = 0.28 *A*_1_ = −0.82; *R*^2^ depicted in the graph).

To look at fusion competence, we calculated the probability of release, that is, how likely a newcomer vesicle was to fuse once it arrived at the membrane, regardless of stimulus. **Figure 6B** shows a cumulative histogram of release probability as a function of sitting time for the same data as in **Figure 7A**. The fusion probability showed time dependence, with no vesicles fusing in response to stimuli within 90 ms of their arrival (as in **Figure 6A**). The cumulative probability graph in **Figure 6B** could be fitted with a single exponential function with time constant of 76.87 ms (insert; adjusted *R*^2^ depicted in the figure). These data indicate that indeed newly arrived vesicles are not yet ready to be released and progressively become more release-ready with time.

Taken together, the distribution of *delays for arrival* (**Figure 3C**), combined with the low probability of fusion of newly arrived vesicles (**Figure 6A-B**) suggest that most vesicles may need 100-350 ms to approach the membrane, depending on departure point, and a subset become release-ready in another 200-500 ms once they have arrived. This would yield recovery times in the order of 300-850 ms, close to the fastest time constant for recovery from paired-pulse depression reported elsewhere (Gomis et al., 1999).

## DISCUSSION

### Unidirectional Flow of Vesicles on the Ribbon

Based on its morphology, the synaptic ribbon has long been hypothesized to be a vesicle-transporting organelle (Bunt, 1971; Gray and Pease, 1971). Our study is consistent with this hypothesis, since the vesicles we observed moved in all cases towards the membrane with no retrograde motion, indicating an extreme directional bias in their movement direction.

An elegant previous study using confocal imaging of sparsely labeled vesicles in the analogous bipolar cell in zebrafish showed a milder directional bias in vesicle movement on the ribbon, with most vesicles moving toward the membrane, but a significant portion moving away (Vaithianathan et al., 2016). While species differences is a formal possibility, we view this as unlikely. We instead consider two, not mutually exclusive, other explanations for this difference: 1) vesicles at distal sites, invisible to TIRF imaging, behave differently than the ones nearest the membrane; 2) The signal-to-noise ratio with confocal imaging prevented precise enough determination of vesicle location in some vesicles, causing some vesicles to appear to go in a retrograde direction.

Our approach makes use of the high signal-to-noise afforded by TIRF microscopy and the labeling of vesicles with many fluorophores to resolve nm-scale movements of individual vesicles. In a previous publication, we estimated a frame-to-frame jitter of 3.8 nm in the z-direction for immobilized vesicles, which may reflect either noise or genuine movement and can be thought of as an upper limit for localization in the z-direction (Zenisek, 2008). The work of Vaithianathan and coworkers tracked individual vesicles, likely containing a single fluorescent molecule (Vaithianathan et al., 2019), at an average resolution of 27 nm, with many docked vesicles showing jitter in localization of greater than 50 nm. This suggests that the approximately 30-40 nm movements we observe here may be obscured in their experiments by the noise for some vesicles.

However, it should be noted that our superior resolution comes at a cost of not being able to observe distal locations on the ribbon. Our TIRF microscope generates an evanescent field with a length constant of approximately 50 nm, allowing us to visualize vesicles at distances up to around 100 nm and perhaps less, depending on the signal-to-noise of the labeling. The ribbon in bipolar cells projects distally into the cell up to 150 nm, making much of the back half invisible to our imaging, indicating that distal vesicles moving in a retrograde direction would not be detected in our studies.

### Vesicle Movement in Response to Stimulus Trains

In this study, we investigated the properties of vesicle transport and docking in bipolar cell ribbon-type presynaptic terminals. We find that docked vesicles are concentrated on the ribbon and that ribbon-associated vesicles move rapidly along the ribbon in response to repetitive stimuli. We also found that the vesicle *delay for departure, transit time* and vesicle speed slowed with repeated stimuli, while distances traveled were not significantly different. Since Ca^2+^ levels lower dramatically with distance from open Ca^2+^ channels (Naraghi and Neher, 1997; Roberts, 1993), Ca^2+^ concentration is expected to drop at more distal locations on the ribbon. Hence, one explanation for this finding would be that the trigger for initiation of movement is Ca^2+^-dependent and that lower Ca^2+^ concentrations at distal sites lead to a slower initiation of movement. Alternatively, distal vesicles may have to wait for vesicles beneath them to move, before they can begin movement. Further experiments will be necessary to distinguish between these two models.

Interestingly, the number of departures mirrored the number of fusion events in the preceding stimulus. Since the bipolar cell L-type Ca^2+^channels are slowly inactivating (Heidelberger and Matthews, 1992; Tachibana et al., 1993), and Ca^2+^ levels return to basal levels over many seconds (Kobayashi et al., 1995; Zenisek et al., 2003; Zenisek and Matthews, 2000), Ca^2+^ levels rise with recurring depolarizations (Kobayashi et al., 1995). If Ca^2+^ were the trigger for movement, one would not expect the initiation of movement and translocation itself to be slowed down with repeated stimulation. Hence, exocytosis and presumably vacancies at the base of the ribbon rather than Ca^2+^ itself appear to be the determinant for the number of vesicles recruited. While our results favor a vacancy model, experiments will be necessary to distinguish between these two models.

### Vesicle Depletion and Depression at Bipolar Cell Synapses

We show here that the number of newcomer vesicles correlated with the number of vesicles released during each stimulus for interstimulus intervals equal to or longer than 250 ms, indicating that vesicle replenishment is not rate-limiting for those interstimulus intervals. For shorter interstimulus intervals, vesicle depletion played a prominent role in synaptic depression.

Although the repopulation of active zones was rapid, newly immobilized vesicles were not immediately available for release. Our results directly show that vesicles move down the synaptic ribbon in response to depolarization and that delivery of vesicles to the membrane is not sufficient to make these vesicles release-ready. Together, the time vesicles need to arrive at the membrane and the time they need to become release-ready might explain the long recovery times from paired-pulse synaptic depression.

### Paired-Pulse Depression and Vesicle Availability

Many synapses exhibit short-term synaptic depression in response to two closely spaced stimuli. Considerable evidence indicates that depression arises from a depletion of the pool of vesicles ready for immediate release and its recovery reflects a refractory period in which new vesicles replenish this pool (Zucker and Regehr, 2002). It is not known, however, to what extent depression reflects the physical absence of a vesicle and whether the physical replenishment of these vesicles reflects recovery from depression. Here, we investigated the relationship between vesicle release, replenishment and synaptic depression in goldfish retinal mixed-input bipolar cells by directly imaging single vesicles using TIRF microscopy. The synaptic terminal of a goldfish mixed-input bipolar cell contains about 30-70 ribbons (Holt et al., 2004; von Gersdorff et al., 1996; Zenisek et al., 2004), each carrying about 100 synaptic vesicles (von Gersdorff et al., 1996). In addition, there are hundreds of thousands of vesicles scattered randomly throughout the cytosol (von Gersdorff et al., 1996).

Based on the work of many investigators, at least three components of exocytosis have been identified: a small rapid component, which is exhausted with a time constant of approximately 1.5 ms (Mennerick and Matthews, 1996; Neves and Lagnado, 1999; Sakaba et al., 1997; von Gersdorff et al., 1998), a slower component that is exhausted with a time constant of about 300 ms (Gomis et al., 1999; Sakaba et al., 1997; von Gersdorff and Matthews, 1994; von Gersdorff et al., 1998) and a sustained component (Lagnado et al., 1996; Rouze and Schwartz, 1998). Similar results have been observed in recordings from mouse and rat bipolar cells (Singer and Diamond, 2006; Zhou et al., 2006). Since the combined size of the fast and the slow component of exocytosis is strikingly similar to the total number of vesicles on the ribbon and the size of the fast component is similar to the number of vesicles at the base of the ribbon, these two components or pools are thought to represent the fusion of all vesicles tethered to the ribbons prior to the stimulus (von Gersdorff et al., 1996), the fast component corresponding to fusion of vesicles at the base of the ribbon. Our results here agree well with this idea: we show that single 30 ms depolarizations cause exocytosis of membrane proximal vesicles that are rapidly replenished within 150-350 ms after the termination of the pulse.

Retinal bipolar cells exhibit paired-pulse depression with a time constant of recovery that depends on stimulus strength (Burrone and Lagnado, 2000; Gomis et al., 1999; Mennerick and Matthews, 1996; Singer and Diamond, 2006; von Gersdorff and Matthews, 1997). One study using membrane capacitance reported that recovery from paired-pulse depression in response to a 20 ms stimulus followed two time constants of 0.64 and 31 s, with about 70% of the vesicles exhibiting the longer time constant (Gomis et al., 1999). Our results show that vesicles removed by exocytosis are replenished in approximately 100-350 ms and need another 200-500 ms to mature, which would explain the fastest time constant found by these authors.

### The Lower Release Probability of Newly Immobilized Vesicles

The data presented here indicate that, although vesicle replenishment is fast, newly immobilized vesicles are not immediately competent for fusion. This results in a decreased release probability of these captured vesicles when compared to vesicles that had been docked for some time. There are some hypotheses that try to explain this phenomenon. In some systems, a positional priming step is required to localize vesicles to Ca^2+^ channels (Hwang et al., 2013). Consequently, the Ca^2+^ concentration would be diminished at the new release sites, leading to an apparent reduction in release probability until channels can find their way to the vesicles. Our results could be explained by such a mechanism. An alternative explanation for this change in release probability is that synaptic vesicles are intrinsically heterogeneous, and that release probability is modulated actively according to the stimulation history of a particular synapse (Burrone and Lagnado, 2000; Wolfel et al., 2007). This possibility is related to the fact that fusion requires vesicles to undergo a maturation or priming process that includes more steps than immobilization itself, bringing vesicles closer to being releasable, yet preventing them from being released spontaneously (Sorensen, 2004).

### Comparison with Conventional Synapses

To help understand the role of the synaptic ribbon in synaptic transmission, it is useful to compare the vesicle dynamics in ribbon-type and non-ribbon type synapses. In response to trains of 30 ms depolarizations, new vesicles moved toward the membrane to re-populate the base of the ribbon within 100-200 ms, which was followed by a period of 200-500 ms before vesicles became fusion competent. Previous studies using prolonged stimuli also showed similar delays between vesicle arrival and exocytosis in salamander rods [90 ms (Chen et al., 2013)] and bipolar cells [200 ms (Zenisek et al., 2000)] and also show fast replenishment.

Striking differences emerge when comparing TIRF measurements at conventional synapses to those we measure here. In hippocampal mossy fiber boutons (Midorikawa and Sakaba, 2017) and in the isolated calyx of Held (Midorikawa and Sakaba, 2015), the physical replacement of vesicles was a comparatively slow process, taking 4-5 seconds, 40 to 50 fold slower than ribbon synapses. Despite the slow replenishment of vesicles to the membrane, recovery from depression is relatively fast via the priming of previously docked vesicles. This priming of pre-docked vesicles occurs over a similar time scale as the maturation of newcomers in our study or in photoreceptors, but in the case of conventional synapses Midorikawa and Sakaba (2015, 2017) found that newcomer vesicles did not become release-ready over the time of their imaging, indicating that the transition from vesicle arrival to a pre-primed state was too slow to measure in conventional synapses. These results indicate that ribbon synapses both re-supply new vesicles and speed up a vesicle priming or pre-priming step. The role of ribbons in catalyzing vesicle maturation is also congruent with experiments where acute photodamage to the ribbon eliminates a release downstream of its ability to bind vesicles(Snellman et al., 2011).

While our results support ribbon synapses as specialists in rapid and efficient delivery of release-ready vesicles, it remains uncertain as to why genetic removal of Ribeye leads to profound effects on ribbon morphology, but mild effects on kinetics of continuous exocytosis (Becker et al., 2018; Jean et al., 2018; Lv et al., 2016; Maxeiner et al., 2016). We suggest three hypotheses to explain the results. 1) that the structure itself is relatively unimportant for vesicle delivery and preparation and that the specialized proteins in ribbon synapses can fulfill the role in the absence of a ribbon. Indeed, while Ribeye is necessary for formation of normal ribbons, it seems dispensable for retaining normal release kinetics in hair cells and mouse bipolar cells. 2) Genetic and/or homeostatic compensation in knockout animals can overcome the loss of the ribbon to return release kinetics to normal, perhaps invoking other mechanisms besides vesicle trafficking. 3) The ribbon’s role in goldfish mixed-input bipolar cells is different than in other cell types. Future experiments will be needed to address these possibilities.

## METHODS

All experiments were approved by the Yale Animal Care and Use Committee and were performed according to the *ARVO Statement for the Use of Animals in Ophthalmic and Visual Research*.

### Cell Preparation

Goldfish bipolar neurons were prepared as previously described (Joselevitch and Zenisek, 2009; Zenisek et al., 2002). Briefly, goldfish (*Carassius auratus*) were decapitated and eyes enucleated and hemisected. Retinas were isolated, cut into four to six pieces and incubated at room temperature for 20 minutes in a low-Ca^2+^ solution designed to stop exocytosis, containing (in mM): 120 NaCl, 0.5 CaCl_2_, 2.5 KCl, 1.0 MgCl_2_, 10 glucose, 10 HEPES, 0.75 EGTA (260 mOsm, pH adjusted to 7.4 with NaOH), plus 1100 units/mL hyaluronidase (type V, Sigma). Next, the pieces of retina were washed in low-Ca^2+^ solution and placed for 30-35 minutes in a digestion medium consisting of low-Ca^2+^ solution with 35 units/mL papain (lyophilized powder; Sigma) and 0.5 mg/mL cysteine. Finally, the pieces of retina were rinsed and placed in low-Ca^2+^ solution in an oxygenated environment at 14 °C until dissociated. Retinas were triturated mechanically with a fire-polished glass Pasteur pipette and plated onto a highly refractive coverslip (*n*_488_ = 1.78; Plan Optik AG, Elsoff, Germany) for recording and imaging. All recordings were performed within 90 minutes of dissociation.

### FM1-43 Loading, Imaging and Data Acquisition

For imaging studies, a selected cell terminal was puffed for 5 s with a solution containing 3-5 μM FM1-43 (Molecular Probes) and (in mM): 2.5 CaCl_2_, 25 KCl, 97.5 NaCl, 1.0 MgCl_2_, 1 Trolox® (6-hydroxy-2,5,7,8-tetramethylchroman-2-carboxylic acid, Sigma) and 10 HEPES (250 mOsm, pH adjusted to 7.4 with NaOH). This solution is expected to depolarize the terminals and stimulate exo- and endocytosis. The neuron was then washed by local superfusion for 30–60 min with the low-Ca^2+^ solution designed to stop exocytosis. In some experiments, 1 mM ADVASEP-7 (Sigma) was added to this washing solution in an effort to reduce background staining (Kay et al., 1999). After the wash, the superfusion was switched to the recording solution, containing (in mM): 120 NaCl, 2.5 CaCl_2_, 2.5 KCl, 1.0 MgCl_2_, 10 glucose, 10 HEPES and 2 glutathione (260 mOsm, pH adjusted to 7.4 with NaOH).

For patch-clamp recordings, 8-12 MΩ thick-walled borosilicate electrodes (BF150-86-10HP, Sutter Instrument Company, Novato, CA) were pulled with a P-97 Brown/Flaming Puller (Sutter Instrument Company, Novato, CA) and filled with a solution containing (in mM): 100 Cs-methanesulphonate, 10 TEACl, 4 MgCl_2_, 10 HEPES, 0.5 EGTA, 10 ATP-Mg, 1 GTP-Li and 1 glutathione (230 mOsm, pH adjusted to 7.2 with CsOH). For synaptic ribbon visualization, the pipette solution also contained 5 μM of a Ribeye-binding peptide (rhodamine + EQTVPVDLSVARDR, m.w. 1997.75), synthesized by the W.M. Keck Facility at Yale University.

Cells were viewed through an inverted microscope (IX70, Olympus) modified for through-the-objective TIRFM (Axelrod, 2001). A 488 nm wavelength beam from a solid-state laser (Coherent, Santa Clara, CA) was applied to image FM1-43 fluorescence, while a 561 nm laser (CV Melles Griot, Carlsbad, CA) was employed to visualize the fluorescence of the Ribeye-binding peptide. Both beams were expanded and focused off-axis onto the back focal plane of a 1.65 NA objective (Apo ×100 O HR, Olympus). Two shutters (Uniblitz, Rochester, NY) placed in the optical paths controlled the illumination. After leaving the objective, light entered immersion oil of high refractive index (*n*_488_ = 1.78, Cargille Labs, Cedar Grove, NJ) and then a coverslip of similar refractive index glass (*n*_488_ = 1.78, Plan Optik AG, Elsoff, Germany). The beam was totally internally reflected at the interface between the glass and the solution or cell, generating an evanescent field with a decay constant of around 50 nm, which allowed us to see about 100 nm within the cell. Fluorescence was recorded using an on-chip amplified CCD camera (Cascade 512B, Photometrics, Tucson, AZ). Image sequences were captured at 33 Hz with MetaMorph Software (Molecular Devices, Downington, PA).

### Imaging Data Analysis

Results were analyzed with MetaMorph and Matlab (Mathworks Inc., Natick, MA). To find the location of docked vesicles and ribbons, images were averaged and fit to a 2D-Gaussian function on an inclined plane. Each individual frame was also fit to a Gaussian function and the standard error in the location of the center was used as an estimate of the error in the determination of the center of the object. For high-resolution localization (**Figure 1D**), vesicles and/or ribbons with an error larger than 20 nm in location were discarded. For other types of analyses, all vesicles within 300 nm of the nearest ribbon were used.

Movies with fusing, captured or moving vesicles were visually selected and square regions centered on the vesicle were excised for analysis using Matlab subroutines. To estimate the timing of vesicle departure and arrival, the fluorescence of vesicles was measured in a 0.6-μm diameter region of interest (**Figure 2B**, black circle) and subtracted from the background fluorescence within a 1.2-μm diameter concentric annulus (**Figure 2B**, grey circle). Fluorescence datapoints were fit with a Boltzmann function of the form (**Figure 2C**, red trace):

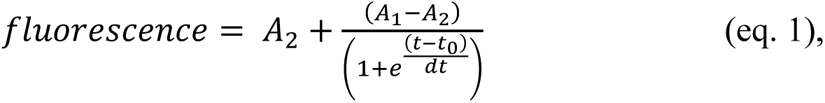

where *A*_2_ (maximal amplitude), *A*_1_ (minimal amplitude), *t*_0_ (function midpoint or half-maximal time) and *dt* are free parameters. The *t* value corresponding to 90% of *A*_2_ was defined as the *arrival time* of newly immobilized vesicles and calculated as:

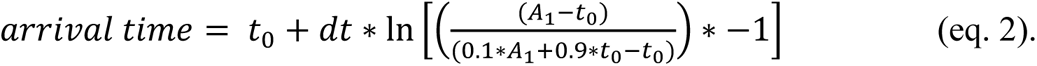

Similarly, the moment vesicles started moving after a voltage stimulus was called *departure time* and defined as the *t* value corresponding to 10% of *A*_2_. It was calculated as:

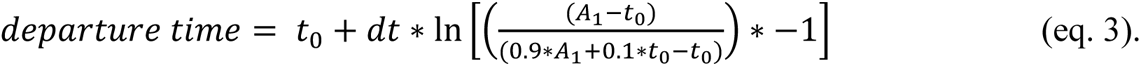

Other forms of analysis for estimating vesicle dynamics are described in the text.

To have an estimate of the distance traveled by a vesicle each time it makes a step towards the membrane (**Figure 2D**), we calculated the vertical displacement for each vesicle (*D*_z_) from the ratio of fluorescence between the start and end of the movement (*A*_2_/*A*_1_). according to a method described elsewhere (Zenisek et al., 2000):

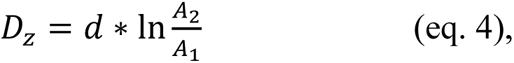

where *d* is the decay constant of the evanescent field (50 nm in our experimental conditions).

Vesicle speed was determined by fitting the fluorescence profile of individual vesicles (or the mean fluorescence profile of all vesicles) to a single exponential followed by an abrupt stop, using a least-squares method for error minimization.

### Statistical Analysis

Statistical tests were performed using Origin Pro 8 software (OriginLab). Data was checked for normality with the Shapiro-Wilk test. Non-normal data was further analyzed with two-tailed Mann-Whitney tests; accordingly, medians are reported in addition to means.

## ACKNOWLEDGMENTS

This work was funded by grants from the National Institute of Health (EY021195, EY003821), the Yale University Vision Core (EY026878) and FAPESP (2010/16469-0 to CJ).

## AUTHOR CONTRIBUTIONS

CJ and DZ designed and performed experiments, analyzed data and wrote the paper.

