## Supplementary material for "Direct Observation Of Vesicle Transport On The Synaptic Ribbon Provides Evidence That Vesicles Are Mobilized And Prepared Rapidly For Release": movie legends

**Supplemental movie 1.** Squares indicate the coordinates of the approach of two synaptic vesicles and fusion of one of them in response to a 30-ms voltage step from -60 to 0 mV. Movie frames were low pass filtered to remove high spatial frequency pixel noise.

**Supplemental movie 2.** Squares indicate the coordinates of the fusion of three resident vesicles and subsequent approach and release of newcomers in response to a 30-ms voltage step from -60 to 0 mV. Movie frames were low pass filtered to remove high spatial frequency pixel noise.

**Supplemental movie 3.** Bipolar cell terminal loaded with FM1-43 and submitted to four consecutive 30-ms depolarizing steps from -60 to 0 mV at 480 ms interstimulus intervals (timing of the stimulus indicated by white circle). The first frame of the movie is the same terminal imaged with the 561 nm laser, showing the location of the labeled ribbons (same as Figure 2C, *Rpep*). Movie frames were low pass filtered to remove high spatial frequency pixel noise.

**Supplemental movie 4.** Average of 5 triggered vesicle approaches, showing that the movement of synaptic vesicles upon stimulation is a localized increase in fluorescence, without lateral spread.
